# The impact of elevated aestivation temperatures on the behaviour of Bogong Moths (*Agrotis infusa*)

**DOI:** 10.1101/2022.10.03.510708

**Authors:** Rose M Lownds, Christopher Turbill, Thomas E White, Kate DL Umbers

## Abstract

Bogong moths are an iconic Australian insect that migrates annually in spring from low elevation locations in southern and eastern Australia to the Australian Alps where they aestivate during summer. As summer ends they make their return journey to the breeding grounds where they mate, lay eggs and die. Given the moth’s extreme behaviour in seeking out cool alpine habitat and with the knowledge that average temperatures at their aestivation sites are rising because of climate change, we assessed whether increased temperatures affect Bogong moth activity during aestivation. Our first hypothesis was that moth activity would be affected by temperature, and we found that moths were more active at higher temperatures, especially during the day, with near-constant activity at 15 °C at all times of day. Our second hypothesis was that moth mass would be different after aestivating at different temperatures for a week due to dehydration or consumption of body energy reserves. We found that moth wet mass loss increased with increasing temperature, but found no difference in dry mass among temperature treatments. Overall, our results suggest that Bogong moth activity will increase with increasing temperatures at their aestivation sites. The impact of warming on the success of individuals to complete their aestivation and journey back to their lowlands to breed should be investigated as a matter of priority to better understand the impact of changes in aestivation behaviour on population dynamics.

## Introduction

Aestivation is a period of behavioural dormancy often facilitated by a down-regulation of physiological processes, similar to hibernation but that it occurs in summer rather than winter (Withers and Cooper 2010). Aestivation is characterised by whole-body metabolic depression, including inhibited digestion and muscular movements (Cowan et al. 2000) and paused growth and development (Masaki 2009), and this behaviour is most common in ectotherms including many insects (Dao et al. 2014, Young et al. 2011). The function of aestivation is to severely depress the organism’s metabolic rate to enable dormancy and prolong survival when in an unfavourable summer environment, e.g. elevated temperature and reduced food and water resources, until conditions improve (Storey and Storey 2012). For example, the mosquito *Anopheles coluzzii* aestivates when there are no viable larval sites during the Sahelian dry summers (Dao et al. 2014), and several African snail species aestivate during periods of drought (Rubaba et al. 2016).

As the climate change drives hotter and drier conditions in many places, species’ abilities to successfully aestivate are becoming increasingly uncertain. Elevated temperatures are likely to disrupt aestivation, particularly in ectotherms where changes in external temperature can directly impact body temperature and metabolism (Young et al. 2011). Increases in metabolism while aestivating may lead to the organism’s energy reserves being depleted more rapidly than the rate at which environmental conditions become more favourable. Furthermore, increased temperatures and subsequently increased metabolic rate may have a compounding effect if organisms become more active while aestivating in warmer temperatures (Lorenz and Gäde 2009).

The impact of climate change induced increases in temperature on aestivation have been observed in several species. Mech et al. (2018) found that elevated aestivation temperatures significantly increased the mortality rates of the insect species hemlock woolly adelgid (*Adelges tsugae*). It was suggested that this was due to factors such as increased heat-induced dessication and elevated metabolic rates (Sussky and Elkington 2015, Neven 2000), though the elevated temperatures may have additionally led to disruption of the insect’s bacterial endosymbionts due to heat shock (Dunbar et al. 2007). Elevated aestivation temperatures can also lead to species decline through habitat loss. A study conducted in south-western Australia of aestivating fish species *Galaxiella nigrostriata* and *Lepidogalaxias salamandroides* found that there has been a total extirpation of these species from 28% and 33% of their historical sites (Ogston et al. 2016). This is thought to be due to altered habitat quality and niche availability as a result of elevated water temperatures caused by global warming (Morrongiello et al. 2011, Ogston et al. 2016).

Aestivating organisms vary in their response to increased temperatures. Some species make no changes and simply pay the costs of temperature changes at the aestivation sites, perhaps because they are unable to move into more favourable environments. Such species may face an increased risk of mortality while dormant as a result of their energy reserves becoming exhausted before their habitat returns to favourable conditions. For example, it was found that when the frog species *Cyclorana alboguttata* experienced elevated temperatures under laboratory conditions during aestivation, its metabolic rate significantly increased contributing to depleted fuel stores and subsequent muscle disuse atrophy (Young et al. 2011). Species that migrate to their aestivation sites may change the time of year or duration spent at the site (Turbill and Prior 2016), or potentially alter arrival and departure times to and from the aestivation sites. Other species may change the location of aestivation, for example several burrowing frog species can mitigate the effects of rising temperatures on their metabolic rate while aestivating by burrowing into cooler ground (Young et al. 2011).

Bogong moths (*Agrotis infusa*) are an iconic Australian insect famous for their annual return migration from the lowlands of south-eastern Australia to the Australian Alps in summer up to a 1000 km away (Warrant et al. 2016). They are of deep cultural value to First Nations Australians, important to alpine ecosystem functioning (Green et al 2020), and a significant pest of wheat and barley (Common 1958). Due to a crash in the number of moths arriving at the alps since 2019 (Green et al 2020) – ~5% of the normal 4 billion expected – Bogongs are now listed by the IUCN as Endangered and currently being assessed for the national legislation, Environmental Protection and Biodiversity Conservation Act. The Australian Alps are warming quickly with the distributions of many Australian mountain species expected to change in the next 25 years (Camac et al. 2020). The Murray murrumbidgee region which includes part of the Kosciuszko National Park has a projected increase in mean temperature of 0.62°C between 2020-39 and 1.94°C between 2060-79 (NSW Interactive climate change projections map 2014), whereas, under the best-case scenario, projected an increase for the Australian Alps are 0.6°C by 2050, and under the worst-case scenario, an increase of 2.9°C (Pickering 2007, Hennessy et al. 2003). There are also projections of reduced rainfall within the Australian Alps by 2090 (Clarke et al. 2019).

Bogong moth aestivation sites are warming and the impact of these rising temperatures are currently unknown. This study aims to determine how Bogong moths respond to increased temperatures while aestivating. There are many important reasons to understand and attempt to mitigate consequences of climate change on Bogong moth aestivation. They are of significant cultural importance to First Nations Australians from across different Countries, who historically, gathered to conduct moth hunts for ceremony and food (Warrant et al. 2016, Flood 1980, Flood 1996). The moths decline represents another fading opportunity for First Nations Australians to reconnect with country and culture. Elevated temperatures in the mountains may already have caused changes in the moth’s arrival and departure times, with moths observed by Caley and Welvaert (2018) to be arriving and leaving nearly a full month earlier than the moths studied by Common in 1954, which arrived at the end of October. Changes in moth arrival times and distribution of aestivation sites may have important impacts on Australia’s alpine ecosystem. For example, some species are largely dependent on moths as a main food source or host. These include the mountain pygmy possum (*Burramys parvus*) and the Bogong obligate symbiont worms *Amphimermis Bogongae* and *Hexamermis cavicola* (Green 2010a, Green 2010b, Common 1954, Welch 1963). Bogong moths are of high ecological importance in the mountains due to their key roles in nutrient and energy transport, particularly in transport of nitrogen and phosphorus (Green 2011).

To understand how Bogong moths respond to increased temperatures while aestivating, we tested two hypotheses. The first hypothesis was that there is a difference in activity level of moths at different temperatures, with the prediction that there would be a higher activity levels in the warmer conditions than cooler conditions. The second hypothesis was that moths lose body mass at different rates in different temperatures, with the prediction that increased temperature leads to an increased rate of body mass loss.

## Materials and Methods

### Study species natural history

Once triggered to begin their annual migration in spring, perhaps by the growing scarcity of edible larval food plants through spring and into summer (Warrant et al. 2016, Common 1954), Bogong moths travel at night for several weeks, with individuals flying up to 1000 kilometres on consecutive nights (Drake and Farrow 1985), to reach their alpine aestivation sites (Common 1954). Once there, they aggregate in small caves and crevices, commonly in granite boulder fields, between approximately 1200-2100 m elevation (Common 1954, Warrant et al. 2016) in astonishing densities of up to 17,000 per m^2^ (Common 1954). Inside the caves and crevices, the moths pack very closely together, in a behaviour known as ‘tiling’, with each moth placing their head and upper body under the wings and abdomen of another moth. The tiling arrangement may decrease their rate of desiccation (Common 1954).

Bogong moths generally aestivate from October to March (Caley and Welvaert 2018) and during aestivation their behaviour is characterised by generally low activity levels and developmental delays, with the moths not sexually maturing until their return to their breeding grounds in autumn (Warrant et al. 2016, Common 1954). While aestivating, the moths do not attempt to mate or feed, though they have been observed ingesting water (Common 1954). Aside from their generally dormant nature while aestivating, some moths display daily periods of activity prior to dawn and after sunset including short flights around their boulders, vibrating their wings and crawling around the rocky outcrops or cave walls (Common, 1954, Warrant et al. 2016). This activity is thought to be triggered by changes in light intensity, which causes pigment migration in the moth’s eyes (Vaduva 2016, Common, 1954), but its function is currently unclear (Karpovich et al. 2009).

### Collection of moth specimens

The Bogong moths included in this study were collected from Charlotte Pass in Kosciuszko National Park, New South Wales, Australia (−36.427891, 148.333679) under NSW NPWS Scientific Permit [SL100835, L. Broome] in early December 2019 using a light trap deployed for 24 hours. The moths were kept together and stored at cool temperatures during transit to Western Sydney University, Hawkesbury Campus (−33.603200, 150.760900).

### Marking and housing moths

160 moths were separated into 16 clear plastic boxes (15cm × 10cm × 5 cm) of ten individuals each, and were initially stored in an incubator at 5°C. CO_2_ was used to make the moths temporarily inactive for weighing and labelling. Each moth was labelled by drawing a number on a forewing using a metallic felt-tip marker with a number from 1-10, and weighed (g) prior to the experiment using a scale sensitive to three decimal places. Three small holes were made in the sides of each box with a soldering iron to allow airflow

### Temperature treatments

We kept moths in one of four different temperature treatments, each of which representative of different climate change scenarios at Charlotte Pass, the origin of the moths used in this study. The lowest temperature 7.5°C was chosen to act as a historical comparison for the likely average temperature of aestivation sites around 50 years ago (Common, 1954). The temperature chosen to most closely represent current aestivating sites for Bogong moths was 10°C (Green, 2010). The two warmer incubators were set at 12.5°C and 15°C to represent the worst case temperature scenario in 50- and 100-years’ time respectively (Pickering, 2007).

### Artificial aestivation set up

Four incubators were held at 7.5°C, 10°C, 12.5°C and 15°C (Memmert IPP750, Panasonic MIR554). The lighting for each incubator was set up using a Jaycar LED aluminium light strip with a Jaycar LED Dimmer Switch and Jaycar Mains Timer with LCD display to simulate dawn/dusk timings for the time of year in Charlotte Pass (dawn: 05:42, dusk: 20:09). An infrared LED torch (Jaycar) was placed on the middle shelf of each incubator to allow observation of the moths at night without disturbing them as most insects are not sensitive to light >550 nm (Shockley Cruz and Lindner, 2011). This meant that in the boxes lit by the infrared light movement could be detected by the cameras at all times of day rather than the non-IR lit shelves whose activity could only be detected during daylight hours. Two out of the four boxes containing moths in each incubator were illuminated by the IR LEDs. A tray of water was placed at the bottom of each incubator to provide a constant high relative humidity. The mean relative humidity measured by miniature temperature/humidity Thermocron iButton loggers ranged from 83.8-93.8% across the incubators for the duration of the experiment.

On the 13^th^ December 2019 the 16 boxes of moths were divided equally and haphazardly among four incubators set at different temperatures (7.5°C, 10°C, 12.5°C and 15°C), with four boxes per incubator and two boxes per incubator shelf. On 18^th^ December 2019 Raspberry Pi motion-capture camera traps (Raspberry Pi Foundation, 2012) were set up on each shelf of each incubator to record moth activity levels (each camera recorded two boxes) 24 hours per day for the duration of the experiment. The camera traps recorded activity levels until the 25^th^ December 2019.

### Question 1: Is there a difference in moth activity over the day when aestivating at temperatures?

The videos produced by the incubator camera traps were downloaded and viewed using VLC media player. One author (RL) screened 7,312 videos in total. In each video the number of moths moving was scored. To ensure the scorer was unaware of the temperature treatments when viewing the videos, the file names originally containing the details of each trap including incubator temperature, time and date were anonymised, and the order of videos randomized, to avoid bias in the scoring. Half of the boxes analysed were recorded at both day and night using the infrared light, and the other half only during the day. Any videos that could not be scored with full confidence due, for example, to large numbers of moths gathering in one place were rescored independently by a different scorer. If there was a difference in the scores of these videos among the two scorers, the video was excluded from further analysis (N=46 videos).

To examine the daily time-course of moth activity under different aestivation temperatures, we fit a generalised additive mixed-model with the proportion of moths moving as our response, temperature (7.5, 10, 12.5, 15) as a fixed effect, and individual smooth terms for time (0-23 hours) at each level of temperature. We specified a binomial error distribution with logit link, and included the identity of each “box” (containing n = 10 moths) nested within experimental day as random effects. For all statistical analyses we used R (v4.1.2; R Core Team, 2021) and the packages ‘mgcv’ (v1.8-40; Woods 2017) for GAMs and ‘emmeans’ for post-hoc tests (v1.7.4-1; Russell 2022) (v1.1–23; Bates et al., 2014). We visually confirmed model assumptions by examining diagnostic plots using the ‘performance’ package (v0.9.0; Ludecke et al. 2021).

### Question 2: Is there a difference in moth mass after aestivating at different temperatures?

#### (a) wet mass

Moths were weighed once in the two days prior to the start of the experiment and twice during the experiment. The initial total mass of moths in each box was calculated and compared to ensure that there were no large differences in total moth mass per box. All moths were taken out of their incubator and weighed on 20^th^ December and 7^th^ January (behavioural data were excluded from weighing days).

To test whether aestivation temperature influences moth wet mass we fit a generalised linear model with the change in each individual’s mass from 20^th^ December to the 7^th^ January as the response, and aestivation temperature (7.5, 10, 12.5, 15) as a fixed effect. We specified a Gaussian error distribution with Identity link function and ran post-hoc contrasts to examine all pairwise differences among temperature treatments.

#### (b) dry mass

To test whether aestivation temperature influences dry mass we fit a generalised linear model with dry moth mass as a response, and aestivation temperature (7.5, 10, 12.5, 15) as a fixed effect. We specified a Gaussian error distribution with identity link, and ran post-hoc contrasts to examine all pairwise differences among temperature treatments.

## Results

### Question 1: Is there a difference in moth activity over the day when aestivating at different temperatures?

There was a total of 7,312 videos triggered by the motion-activated camera traps and scored for moth activity. As cameras were trained on two boxes of moths each, each video was watched twice, once for each box, a total of 14624 views. Videos were screened for the presence and absence of movement, and no movement was present in 9,012 views with the motion sensing cameras which we cannot explain. These were discarded leaving a total 5,612 views of boxes with moth activity. Thus only boxes in which any moths were moving were scored for number of moths moving, and all others were scored as no moths moving.

Our generalised additive models identified strong patterns in moth activity over the 24 hour day/night cycle, which varied between aestivation temperatures (Table 1). At 7.5°C and 10°C, peak moth activity was seen immediately pre-dawn (ca. 0500 hours) and post-dusk (ca. 2100 hours; Fig. 1), with minimal activity in daylight hours. These bimodal peaks in activity reduced fractionally at 12.5°C, before being near-completely eliminated at aestivation temperatures of 15°C. Moth activity during daylight was markedly elevated at 15°C, with only minimal increases to activity evident in the pre-dawn and post-dusk hours that characterise their peak activity periods at 7.5-12.5°C (Fig. 1).

**Table 1:** Parameter estimates and test statistics for a generalised additive mixed model examining the effects of aestivation temperatures on Bogong moth activity levels. Temperature (temp) was included as a fixed effect, and separate smooth terms were estimated for time (hour 0-23) at each temperature. Experimental box nested within experimental day were specified as random effects. Note that temperature waw treated as an ordered factor, such that reported statistics represent tests of differences in smooth terms from the reference level of 7.5°C. Model R^2^ = 0.229.

| *Parameter* | $\beta$ | *SE* | *t* | *p* |
| --- | --- | --- | --- | --- |
| *Intercept* | -0.696 | 0.179 | -3.90 | < **0.001** |
| *temp. 10* | 0.139 | 0.251 | 0.553 | 0.580 |
| *temp. 12.5* | -0.366 | 0.243 | -1.504 | 0.1327 |
| *temp. 15* | -0.561 | 0.259 | -2.160 | **0.031** |

| *Smooth terms* | *Effective d.f.* | *Ref. d.f.* | *F* | *p* |
| --- | --- | --- | --- | --- |
| *s(hour) \* temp. 7.5* | 6.866 | 6.866 | 34.85 | < **0.001** |
| *s(hour) \* temp 10* | 6.494 | 6.494 | 17.18 | < **0.001** |
| *s(hour) \* temp 12.5* | 8.262 | 8.262 | 23.21 | < **0.001** |
| *s(hour) \* temp 15* | 7.614 | 7.614 | 22.46 | < **0.001** |

**Fig 1:**
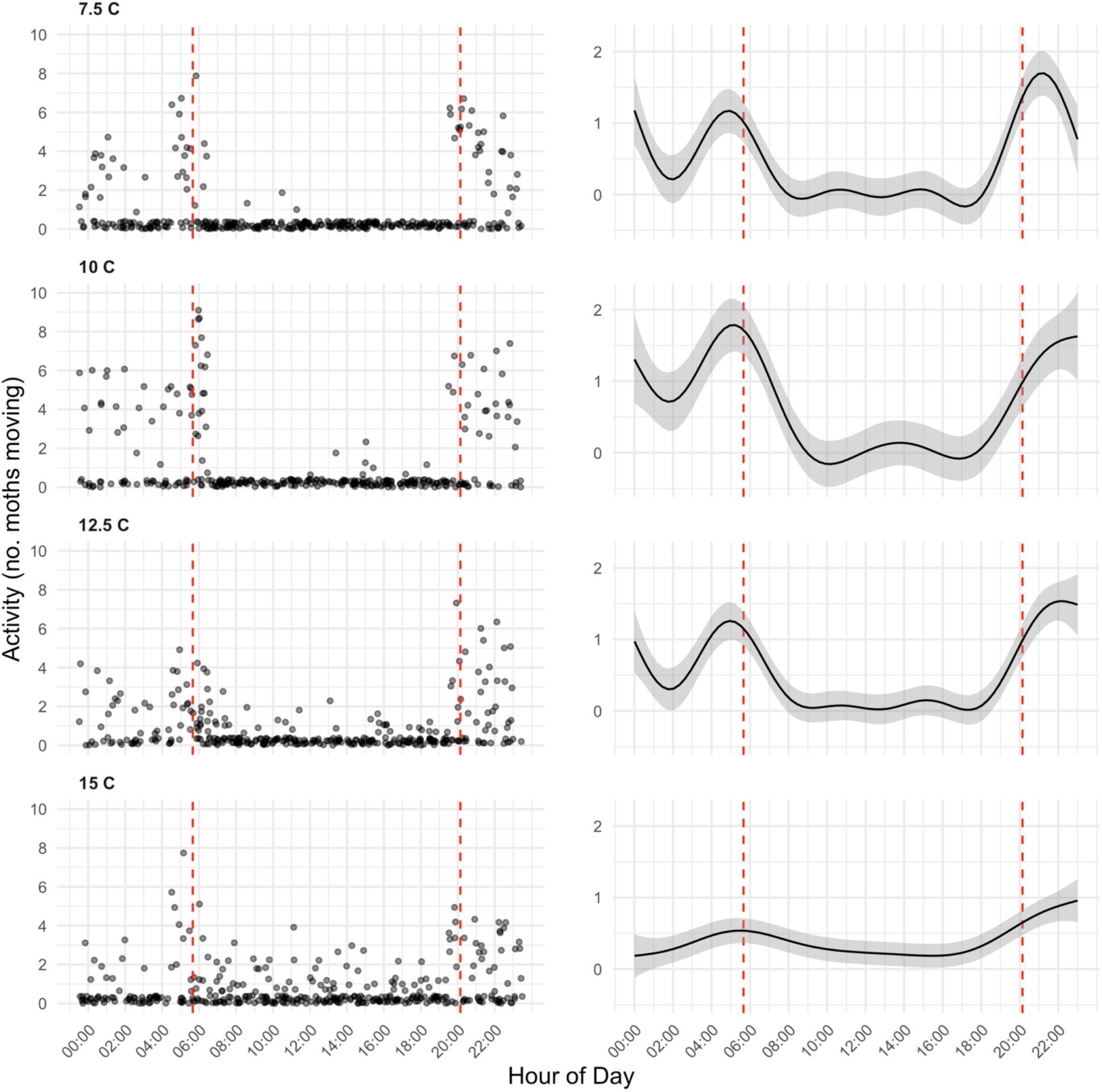
The activity of Bogong moths over 24 hours following aestivation at four different temperatures. Shown are raw data (left panels), denoting the number of moths moving at each timepoint (out of 10 moths total per box, see methods for full sample size & replicate details). And generalised additive mixed-model fits (± 95% CIs, right panels) to these data. Red vertical lines indicate the experimental shift from day to night.

### Question 2: Is there a difference in moth mass after aestivating at different temperatures?

#### (a) wet mass

We identified a moderate to strong effect of aestivation temperature on the loss of wet mass (Fig. 2a). The greatest decline in mass was at 15°C (0.045 ± 0.019g), followed by 12.5 °C (0.038 ± 0.170g), 10 °C (0.032 ± 0.012g), and 7.5 °C (0.023 ± 010g), respectively (see Table 1 and Table S1 for full numerical results).

**Fig 2:**
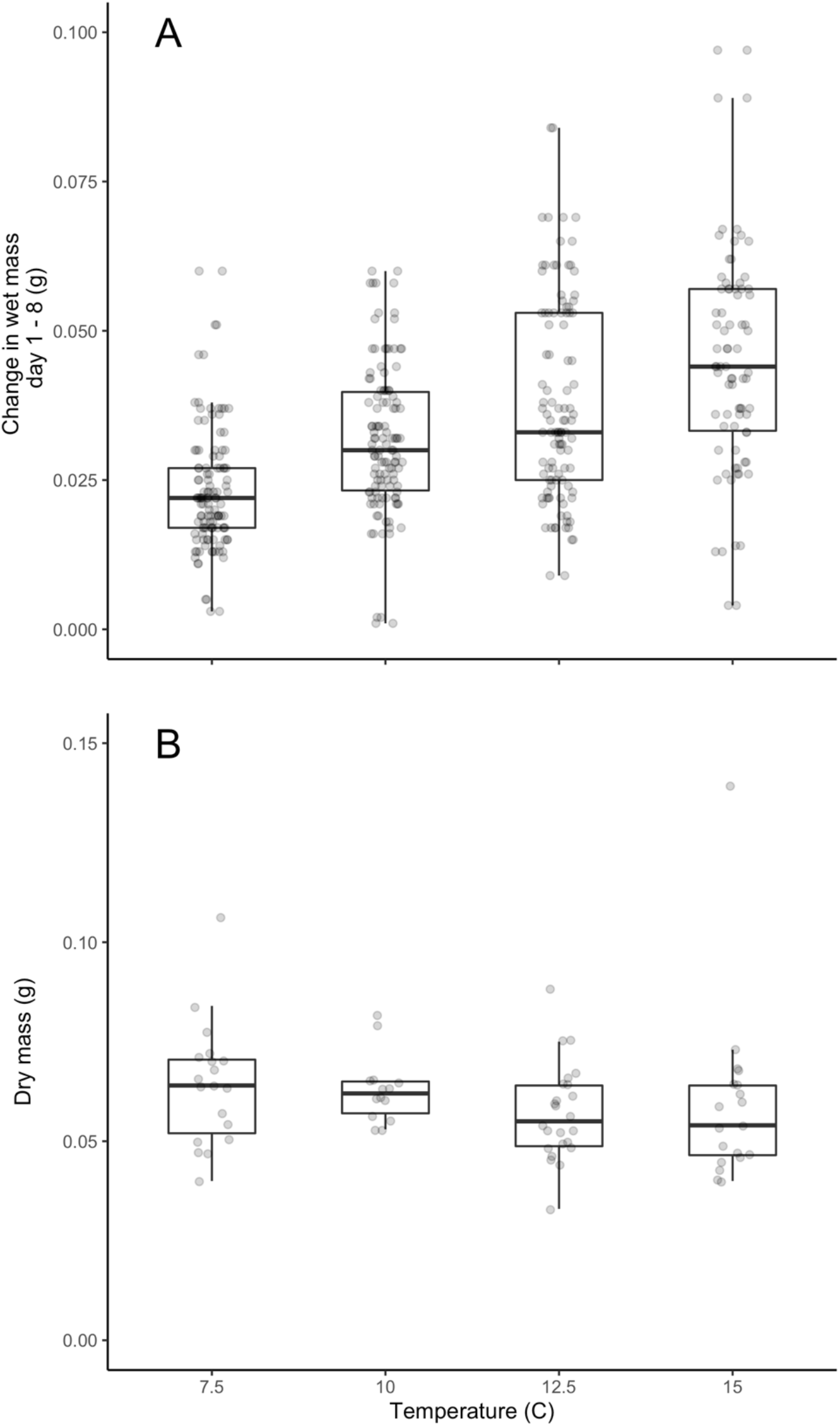
The effect(s) of aestivation temperature on (a) the change in wet mass (day 1 – day 8) and dry mass (b). All Tukey-adjusted pairwise contrasts of wet mass change (a) were statistically significant at α = 0.05 (Table S1), while no measures of dry mass (b) differed between temperature levels (Table S2).

#### (b) dry mass

We found no overall effect of aestivation temperature on the dry mass (Fig. 2b). This held true among all post-hoc pairwise contrasts between temperatures, which did not differ from one another (Table 2, Table S2).

**Table 2:** Parameter estimates and test statistics for a generalised linear model examining the change in Bogong moth wet mass between experimental day 1 and 8, across each of four aestivation temperatures. The difference in mass (day 1 - 8) was specified as a response with temperature as the single fixed effect, and we specified a Gaussian error distribution with identity link function. Model R^2^ = 0.224.

| *Fixed Effects* | $\beta$ | *SE* | *t* | *p* |
| --- | --- | --- | --- | --- |
| *Intercept* | 0.023 | 0.001 | 17.375 | < 0.001 |
| *temp. 10* | 0.008 | 0.002 | 4.530 | < 0.001 |
| *temp. 12.5* | 0.015 | 0.002 | 7.592 | < 0.001 |
| *temp 15* | 0.022 | 0.002 | 10.279 | < 0.001 |

## Discussion

The pattern of daily moth activity differed significantly across the four temperatures. The strong depression in activity evident during daylight hours in the cooler treatments was lost in the warmer temperatures. Bogong moth aestivation behaviour was disrupted at 12.5°C and lost at 15°C. Moths kept at 15°C also had the greatest reduction in wet mass over experiment, but there was no difference in dry mass among temperature treatments at the end of the experiment. Moths seem to dehydrate more rapidly at 15°C than at cooler temperatures. Overall, our results suggest that climate change induced warming of Bogong moth aestivation sites will cause increased moth activity, leading to a reduction in body condition and decreased survival over aestivation. The impact of increased activity during aestivation on moth survival and reproduction (fitness) should be the focus of future investigations.

In our experiment, moths showed peaks of activity at dawn and dusk which matches field observations of moths flying out of their boulder fields in large numbers at dawn and dusk during aestivation (Warrant et al. 2016 and Common 1954). This suggests that broadly, moths were behaving normally in our experimental enclosures. The higher activity levels in the warmer incubators were likely due to increased metabolic rate caused by the higher temperatures (Clarke and Fraser 2004, Deutsch et al. 2018, Neven 2000), which makes greater amounts of energy available for movement (Lorenz and Gäde 2009) perhaps to search for cooler locations.

Increased metabolic rate may impact moth survival. As mentioned earlier, Young et al. (2011) suggested that increases in metabolic rate during aestivation occurring under elevated temperature conditions may impact survival in frog species *C. alboguttata* during aestivation, as a result of the greater energetic cost of physiological functions. This has also been observed in the aestivating insect species hemlock woolly adelgrid (*A. tsugae*) where warmer forests had a higher rate of fatality among the adelgrids while they were dormant (Sussky and Elkington, 2015). If greater moth activity levels and metabolic rate are correlated with higher mortality rates during aestivation as a result of exhausting energy reserves before the organism’s environment has become more favourable, it could reduce Bogong moth abundance change and subsequently alter food chains.

Moth wet mass decreased more rapidly at the higher temperatures, but dry mass did not differ, suggesting that the loss of mass at the higher temperatures was caused by dehydration rather than loss of body fat. Bogong moths have been observed to ingest water on occasion during aestivation in the wild, and so putting a small water source within box of moths may allow drinking. It is possible that over a longer time period fat reserves would be used up during movements and flight at higher temperatures faster than at cooler temperatures. Common (1954) reported decreases in fat content in both male and female Bogong moths between December and January 1951-2 and suggested that it may be partly as a result of moth metabolism during aestivation. Green (2011) measured the mean dry mass of 150 dead moths collected from the mountains as being 0.155g, while the mean dry mass was 0.060g. This supports the idea that the much lower dry masses from this study may be from a combination of increased levels of desiccation and higher activity levels in the incubators than observed in nature, but these comparisons must be investigated.

Future directions following on from this research could include further studying the effects of different temperatures on Bogong moths by measuring their metabolic rate directly by closed-box respirometry where the moth’s gaseous exchange can be directly measured. It would also be useful to determine the critical thermal limits of Bogong moths to further understand the implications of climate change. Male and female moths could also be separated for the duration of temperature studies to determine whether they show different temperature preferences because other insects, *Drosophila melanogaster* for example, show different temperature preferences depending on sex in a similar thermal gradient study (Rajpurohit and Schmidt 2016). Bogongs could be dissected to determine whether they were sexually mature while aestivating at different temperatures, as while studies by Common (1954) found that moths do not become sexually mature during their time in the mountains there were two instances of mating observed in this present study alongside egg laying and hatching. Additionally, it would also be interesting to compare the effects of different temperatures on Bogong moths both from aestivating sites in the mountains and from moths which seemingly do not make a yearly migration (Warrant et al. 2016). These moths are thought to remain in the milder areas of southern New South Wales and Australian Capital Territory (ACT), and may show very different temperature preferences and summer activity levels.

In summary, higher moth activity levels seen in the warmer temperature treatments provide support for an affect of warming from climate change on aestivation behaviour. Increased temperatures may affect the moth’s survival and how any changes in Bogong moth population densities may affect their wider alpine ecosystem. These results may be of importance in aestivation studies for species worldwide as in the event of substantial climate change higher global temperatures may lead to increased metabolic rates and activity levels particularly in dormant ectotherms, with potential increases in mortality rate of aestivating species (Young et al. 2011).

**Table 3:**
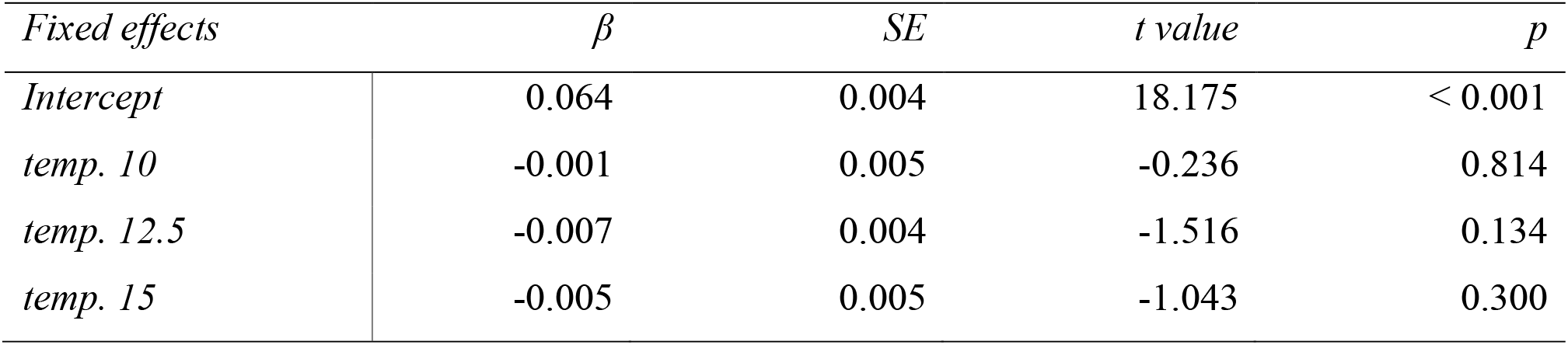
Parameters estimates and test statistics for a generalised linear model examining the dry mass of moths between temperature treatments at the conclusion of the experiment. Model R^2^ = 0.038.

## Acknowledgments

We thank Linda Broome for collecting the moths and Eric Warrant for advice on moth husbandry. KU thanks her family for their continued support, and TEW thanks Liz Mulvenna and Cormac White for the same.

